# It’s the Sound, not the Pulse: Peripheral Magnetic Stimulation Reduces Central Sensitization through Auditory Modulatory Effects

**DOI:** 10.1101/2024.07.30.605813

**Authors:** Spencer S. Abssy, Natalie R. Osborne, Evgeny E. Osokin, Rossi Tomin, Liat Honigman, James S. Khan, Nathaniel W. De Vera, Andrew Furman, Ali Mazaheri, David A. Seminowicz, Massieh Moayedi

**Author notes:** **To whom correspondence should be sent:** Dr. Massieh Moayedi. Co-first authors (equal contributions) (Spencer S. Abssy, Natalie R. Osborne, Evgeny E. Osokin).

## Abstract

Repetitive peripheral magnetic stimulation (rPMS) is a non-pharmacological, non-invasive analgesic modality with limited side effects. However, there is a paucity of controlled studies demonstrating its efficacy compared to existing pain management tools. Here, in an initial sample of 100 healthy participants (age 18-40), we compared the analgesic efficacy of two rPMS stimulation protocols—continuous theta burst stimulation (ctbPMS) and intermittent TBS (itbPMS)—against transcutaneous electric nerve stimulation (TENS), a peripheral stimulation technique that is commonly used for pain management. We also included a sham rPMS protocol where participants heard the sound of rPMS stimulation while the coil was placed over their arm, but received no peripheral stimulation. We hypothesized that itbPMS and ctbPMS—but not sham—would reduce pain intensity, pain unpleasantness, and secondary hyperalgesia evoked by a phasic heat pain (PHP) paradigm on the volar forearm with similar efficacy to TENS. Neither rPMS nor TENS reduced reported pain intensity or unpleasantness (p>0.25). However, ctbPMS and itbPMS significantly reduced the area of secondary hyperalgesia, whereas TENS did not (F_3,96_= 4.828, p= 0.004). Unexpectedly, sham rPMS, which involved auditory but no peripheral nerve stimulation, also significantly reduced secondary hyperalgesia compared to TENS. We performed a second study (n=32) to investigate auditory contributions to rPMS analgesia. Masking the rPMS stimulation sound with pink noise eliminated its analgesic effect on secondary hyperalgesia (p=0.5). This is the first study to show that the analgesic properties of rPMS in acute experimental pain may be largely attributed to its auditory component rather than peripheral nerve stimulation.

## Introduction

Chronic pain affects between 10 and 50% of the global adult population [59]. Current pain management approaches have significant limitations and there are no interventions that are effective for all patients [47]. Complex procedures such as nerve blocks and implantable neuromodulation devices can reduce chronic pain in some individuals, but these interventions are invasive, costly, and associated with significant side-effects [46]. Non-invasive brain stimulation therapies, such as transcranial magnetic stimulation (TMS), have heterogeneous outcomes and variable long-term efficacy [7]. Pharmacological treatments can be associated with significant side-effects and reduced efficacy over time [2,43]. Transcutaneous electrical nerve stimulation (TENS) is a common modality for non-invasive treatment of musculoskeletal pain, given its low cost, and the low effort required for use [53]. TENS, however, can only reach superficial tissues [4]. Therefore, there is limited efficacy in existing pain management tools. There is thus a clear need for novel effective pain management tools.

A novel approach of targeting peripheral tissues with focal magnetic pulses—i.e., peripheral magnetic stimulation (PMS)—is becoming an increasingly popular pain management tool [28,34], largely due to its non-invasive nature and tolerability to physical sensations elicited by PMS [28]. Unlike TENS, PMS can reach deep tissues of the body. In particular, repetitive PMS (rPMS) with theta-burst stimulation (TBS) is of particular interest. Theta oscillations are between 4 and 8Hz, and TBS is delivered at 5Hz. The TBS model was developed to mimic hippocampal LTP/LTD firing patterns [25], and theta rhythms play a key role in memory functions [10,20,23], but their role in pain is less well understood.

A case study by our group reported that repetitive PMS (rPMS) significantly reduced clinical pain in a patient with intractable glossopharyngeal neuralgia [29]. Another study showed that rPMS can modestly reduce pain and improve functional recovery in patients with acute low back pain, compared to a sham rPMS protocol [35]. Moreover, rPMS outperformed sham treatment in patients with low back pain in functional improvement [35].

While rPMS shows promise, there haven’t been good clinical studies with proper controls [38]. This is essential given that rPMS emits rhythmic auditory stimulation, which has been associated with neural entrainment. Indeed, evidence indicates that neural entrainment by rhythmic auditory stimuli can be analgesic [3,13]. Put another way, it is unknown whether the effects of rPMS are driven by the somatic stimulation of peripheral nerves, or by auditory entrainment.

Here, we explore whether rPMS can significantly reduce pain evoked by a phasic experimental heat pain model. Our first aim is to compare whether different rPMS stimulation parameters can reduce pain intensity, unpleasantness, and the area of secondary hyperalgesia, compared to TENS and a sham condition. To account for potential confounding factors, we also conduct a control study to assess the impact of the sounds emitted by the magnetic stimulation device during TBS on rPMS analgesia. Additionally, we examine whether there are sex differences in rPMS analgesia by performing sex-disaggregated analyses

## Methods

### Sample Size Determination

Based on a pilot study with a sample size of n=5, we showed that intermittent theta burst PMS (itbPMS; see below for stimulation parameters) reduces pain intensity, pain unpleasantness and secondary hyperalgesia with effect sizes of Cohen’s *f*= 0.63, 0.81 and 1.3, respectively. Considering the smallest of these effect sizes, the estimated sample size required to achieve 80% power with an alpha=0.0167 (corrected for 3 outcome measures, with a non-sphericity correction of 0=0.87, and a correlation amongst representative measures of R^2^=.39) is 44 participants, using G*Power (v3.1.9.6, Heinrich-Heine-Universität Düsseldorf, Düsseldorf, Germany). Therefore, we need a minimum of 44 participants across 4 intervention arms who complete the study. Furthermore, to perform disaggregated analyses to assess sex differences we doubled the minimum required sample size. We oversampled to ensure we met a minimum of 88 participants for the study, and to account for individuals lost to follow-up or those who withdraw from the study.

### Participants

#### Study 1

A total of 123 healthy participants aged between 18-40 years agreed to participate in the study approved by the University of Toronto Human Research Ethics Board (Protocol# 41856). Exclusion criteria were (1) self-report of neurological, psychiatric or rheumatological disease, including a history of seizures; (2) history of chronic pain; (3) history of chronic illness; (4) clinically significant scores on the Beck’s Depression Inventory (>21), which indicates moderate depression or indication of suicidal ideation (>1 on item 9); (5) <28 on Mini Mental State Exam (6) pregnancy and breastfeeding; (7) heat pain ratings >50/100 for multiple temperatures <40°C (which indicates hypersensitivity); (8) consistent heat pain rating <20/100 for 48°C (which indicates hyposensitivity); (9) inconsistent ratings during the search protocol (i.e., vastly different ratings (>20/100 difference) for the same stimulus intensity, or ratings that do not show increases with increased stimulus intensity); (10) pacemaker or any contraindication to receiving magnetic stimulation; (11) use of medication known to affect pain sensitivity (e.g., NSAIDS, anxiolytic, antispastic, acetaminophen) within 24 hours of testing;(12) visible scarring on the volar forearm testing area; and (13) previous experiences with TMS or PMS. Of these 123 participants, 23 were excluded due to hyposensitivity (n=8), scored >21 on the BDI or reported suicidal ideation (n=3), withdrew (n=2), scars on stimulation site (n=3), inconsistent pain ratings in the pain calibration stage (n=2; see below), missing data (n=2), lost to follow-up (n=2), and hypersensitivity (n=1). The final sample included in this study comprised 100 healthy participants. Figure S1 provides a CONSORT chart of participant recruitment and intervention arm allocation.

#### Study 2

Thirty-two healthy participants aged between 18-40 consented to procedures approved by the University of Toronto Human Research Ethics Board (Protocol# 41856) and were screened for study eligibility, based on the same exclusion criteria as Study 1. Of these, twenty-four participants (12 males and 12 females) met inclusion criteria and were included in the final sample (see Figure S2).

### Experimental design

#### Study 1

Participants underwent two experimental sessions on different days, at least 48 hours apart: a stimulation session and a control session, counterbalanced across subjects (see Figure 2). In the first session, after consenting to procedures, participants were first screened for BDI and MMSE exclusion criteria. Next, they completed the Toronto Academic Health Science Network (TAHSN) demographics questionnaire, and the State-Trait Anxiety Inventory (STAI) [48], as evidence suggests that anxiety can affect pain sensitivity. Participants were then pseudo-randomly assigned to an arm of the study.

**Figure 1.**
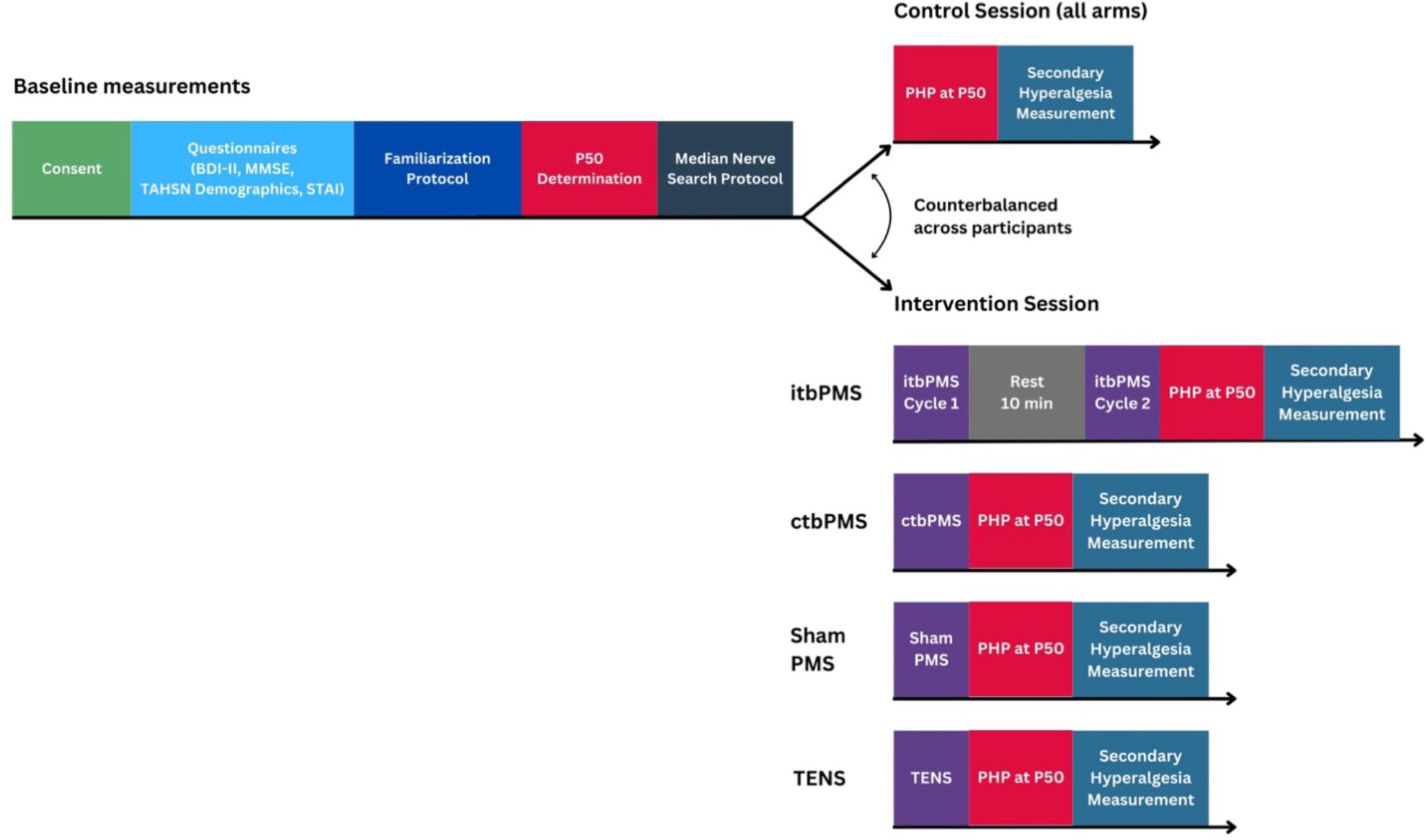
Summary of experimental procedures for each intervention arm in Study 1. Participants attended two sessions: a control session and a stimulation session (arms: itbPMS, ctbPMS, Sham, TENS), separated at least 48 hours. The order of sessions was counterbalanced across participants.

**Figure 2.**
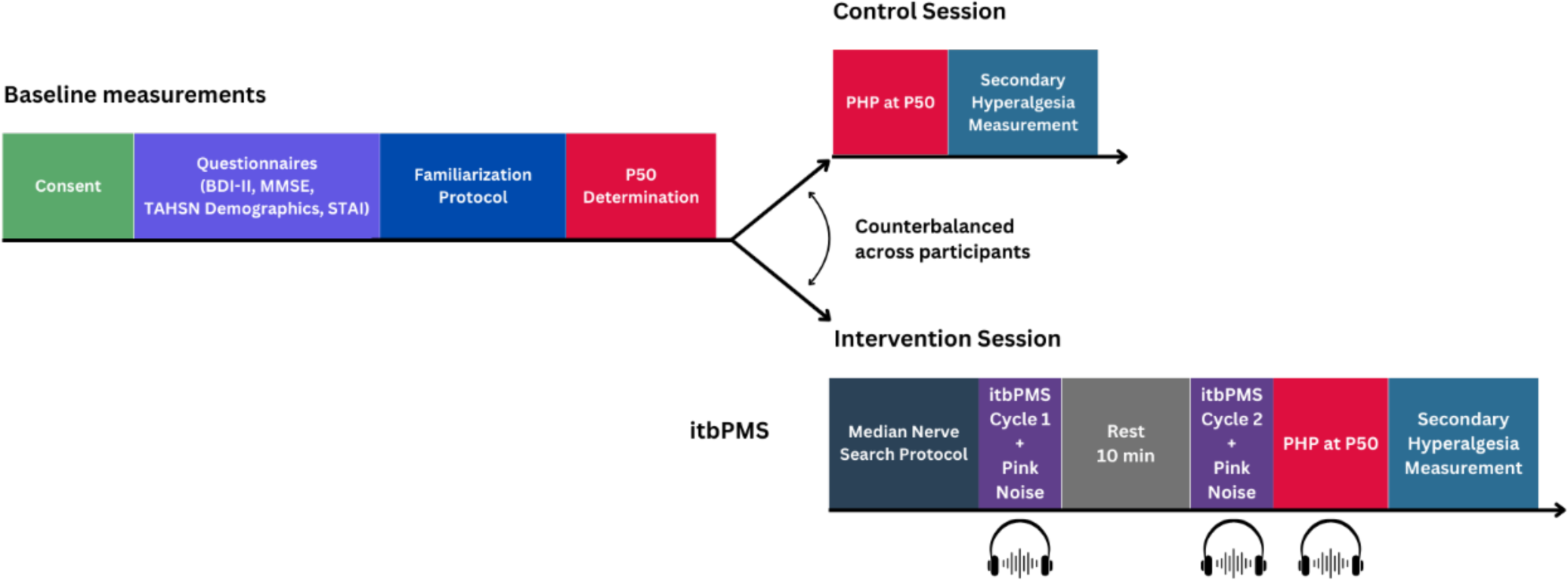
Summary of experimental procedures for the stimulation session in Study 2. Participants attended two sessions: a control session and a stimulation session (itbPMS + pink noise), separated by at least 48 hours. The order of sessions was counterbalanced across participants.

Participants then underwent two further procedures: a familiarization protocol and a search protocol to identify a stimulus intensity that would elicit a pain intensity rating of 50/100 (P_50_). Participants then had an EEG cap mounted, and a 5-minute resting EEG recording was taken. Recording was maintained throughout the experiment with markers for each part of the study (note: we report the EEG data collection for completeness, but these are not analyzed as part of this study and will not be further discussed). Next, the location of the median nerve on the volar forearm was identified for all experiments using a search protocol with the magnetic stimulation coil, and the motor threshold was identified. Depending on the session type (control or stimulation), the participant either received PHP at P_50,_ or a stimulation on the volar forearm over the median nerve territory followed by a PHP at P_50_, respectively. Finally, we measured the area of secondary hyperalgesia elicited by PHP. We then collected another 5-minute resting state EEG recording (not discussed).

*Stimulation Randomization Procedure.* We used a pseudo-randomized approach to assign participants to one of four stimulation groups (itbPMS, continuous theta burst PMS (ctbPMS), TENS, and sham ctbPMS). The order of the control and stimulation sessions were counterbalanced across participants.

#### Heat Stimuli

A thermal stimulation device (QSTLab, Strasbourg, France) with a T11 probe was used to deliver heat stimuli to the volar forearm of participants. Four of the five plates were used to deliver heat over a 3x3cm area.

##### Familiarization

Familiarization consisted of three 7s heat stimuli at three different temperatures (45°C, 43°C, 47°C) with a 15s interstimulus interval. The ramp rate was set at 7°C/s from a baseline of 32°C, to deliver the target temperature for 5.5s before returning to baseline at 7°C/s. This familiarization protocol allowed the experimenter to assess exclusion/inclusion based on pain sensitivity, and familiarized participants to heat stimuli in the noxious range. After each stimulus, participants provided pain intensity and pain unpleasantness ratings using a verbal numerical rating scale (vNRS). The anchors for the pain intensity vNRS were: 0 - “no pain”, and 100 - “worst pain imaginable.” The anchors for the pain unpleasantness vNRS were: 0 - “not unpleasant” and 100 - “most unpleasant imaginable.” Participants were instructed to rate both pain intensity and unpleasantness upon hearing a verbal cue (“Rate Now”) from the experimenter.

##### Pain 50

We individually calibrated PHP stimuli to 50/100 on the vNRS for each participant. To do so, participants received various temperatures (39°C, 41°C, 43°C, 45°C, 47°C). The ramp rate was set at 2.5°C/s from a baseline of 32°C, and the stimulus was held at target temperature for 20s before returning to baseline at 2.5°C/s, with an interstimulus interval of 10s. Each stimulus (except 47°C) was presented twice, and stimuli were presented in a pseudorandomized order with a rectangular distribution. A sham stimulation was also included (at a baseline of 32°C) to ensure that participants were providing true ratings that reflected the stimuli. Participants provided a pain intensity and pain unpleasantness rating after each stimulus. Stimuli that elicited a rating >80/100 were not repeated to minimize injury. A temperature that elicited 50/100 was used for P_50_, and was confirmed with a subsequent stimulus.

##### PHP Protocol

The PHP protocol comprised of five stimuli at P_50_ presented for 40s with a 20s inter-stimulus interval on the non-dominant volar forearm, as previously done [18]. The stimulus had a ramp rate of 7°C/s and was held at the target temperature for 40s. Pain unpleasantness and pain intensity rating were acquired every 10s during each stimulus.

To minimize habituation during familiarization and P_50_ protocols, the thermal probe was moved to a new stimulation site on the dominant volar arm. If at any point participants found the pain from the thermal stimuli intolerable, they were instructed to say “stop” and the thermal probe was immediately removed from the arm.

#### Secondary Hyperalgesia measurement

Secondary hyperalgesia measurement was conducted on the basis of previously published work [40]. Secondary hyperalgesia is defined as the area outside the primary site of stimulation where a noxious mechanical stimulus is felt as hyperalgesic. Punctate mechanical stimuli (256mN PinPrick Stimulator, MRC Systems, Heidelberg, Germany) were used to delineate the border of secondary hyperalgesia. To do so, mechanical stimuli were delivered to the volar forearm along eight orthogonal trajectories radiating from the stimulation site. The stimuli started 6 cm outside the hyperalgesic region and moved toward the stimulation site at 0.5cm increments, until the participant reported an increase in pain sensation. Participants were instructed to report the moment they felt an increase in the sensation of the pinprick. Once they reported the change in sensation, the border was marked, and the procedure was started again along a different trajectory. Trajectories were marked on a digital map, and the area of secondary hyperalgesia was measured in mm^2^ using imageJ [45].

#### Stimulation

##### Identification of the Motor Threshold

The motor threshold of the median nerve was identified once a muscle twitch was observed in the innervation territory of the median nerve (e.g., a thumb, middle and index finger twitch). The stimulation intensity was then lowered to find the motor threshold; i.e., when the twitch was barely perceptible. We purposefully used a crude approach for determining the motor threshold to improve translation of the rPMS paradigm/modality to the clinic in future studies.

##### rPMS Stimulation

rPMS was delivered to the region of the non-dominant volar forearm over the median nerve using a magnetic stimulator (The Magstim Rapid 2+1. Double 70mm AirFilm® Coil (AFC), West Wales, UK). All stimulations were performed at 5% below the motor threshold. The rPMS protocols used in this study comprised of TBS, which refers to a triplet of stimuli presented at 50Hz, repeated at a burst frequency of 5Hz. Two different types of rPMS were used; itbPMS (200 bursts, 1200 pulses total, 41.6 sec cycle time, 2 cycles total, 10 minutes apart) and ctbPMS (500 bursts, 1500 pulses total, 101.0 sec cycle time, 1 cycle total).

##### Transcutaneous Electrical Nerve Stimulation

TENS was applied to the volar forearm over the median nerve using a high voltage constant current stimulator (Digitimer DS7A, Letchworth Garden City, UK) with small, wired hydrogel electrodes (1 x 1 cm; Electrode Store, Impulse Medical Technologies, Buckley, WA). Electrodes were placed along the proximal-distal axis along the median nerve on the volar forearm, 1.5 cm apart. A train of stimulations was presented using a train delay generator (Digitimer DG2A, Letchworth Garden City, UK). The stimulation intensity was determined by slowly increasing the intensity until the participant reported a painful sensation. Next, the intensity was reduced by 2mA to ensure an intense but non-painful stimulation. The TENS protocol lasted 90s using tonic pulses, with the following parameters: 40Hz, 200μs pulse width and the intensity was calibrated to the individual (range: 6-14 mA).

##### Sham Stimulation

This arm of the study exclusively included participants who were naïve to TMS and rPMS. A concealed speaker was installed in the room, behind the magnetic stimulation device. Auditory ctbPMS stimuli were pre-recorded and the volume and the type of sound resembled that of the rPMS stimulation. The coil was placed over the median nerve on the non-dominant volar forearm, and the magnetic strength was set to 0. Participants did not receive magnetic stimulation but did hear the sound emitted by the magnetic device during a typical ctbPMS stimulation.

#### Study 2

Participants underwent two experimental sessions on different days, at least 48 hours apart: an itbPMS stimulation session with pink noise and a control session where no stimuli were delivered, the order of which was counterbalanced across subjects (see Figure 2). Study 2 follows a similar design as study 1 except that EEG was not collected. In the first session, participants completed the BDI and MMSE for screening purposes. Next, they completed the TAHSN demographics questionnaire, and the STAI. Participants then underwent familiarization and the P_50_ search protocol to determine PHP stimulus intensity. Next, in the stimulation session only, participants underwent median nerve localization on the volar forearm using a search protocol with the magnetic stimulation coil, and the motor threshold was identified. Depending on the session (control or stimulation) participants then underwent familiarization and either received itbPMS stimulation on the volar forearm over the median nerve territory followed by a PHP at P_50_, or simply received a PHP at P_50_. Pain intensity and pain unpleasantness ratings were acquired every 10s during each PHP stimulus. During the itbPMS stimulation and PHP participants heard pink noise through Bose Quiet Comfort II headphones (Framingham, MA). Note that noise cancellation was not turned on. Pink noise is characterized by frequencies in the 40-60Hz range and was played off a YouTube video (https://www.youtube.com/watch?v=8SHf6wmX5MU&t=343s; Google, Inc, San Bruno, CA). The intensity of the pink noise was adjusted to participants’ comfort while still loud enough to mask the sounds of rPMS delivery. Finally, we measured the area of secondary hyperalgesia elicited by PHP.

### Statistical Analyses

#### Study 1

##### Questionnaires

We performed a one-way ANOVA with Welch’s test for non-equal variances to compare STAI-T, STAI-S, and BDI scores between intervention arms, as differences in these scores could affect pain sensitivity. Significance was set at p<0.05. In cases where there were differences across intervention arms, the questionnaire measure was included in the models comparing primary outcomes as nuisance covariates.

##### Differences between intervention arms

All statistical analyses were performed in SPSS (v.28, IBM, Armonk, NY). There were three key outcome measures in the study: (1) change in pain intensity, (2) change in pain unpleasantness and (3) change in secondary hyperalgesia. These change measures are calculated as the stimulation session minus the control session.

We performed three non-parametric ANCOVAs, Quade’s test, each with a primary outcome measure (change in pain intensity, change in pain unpleasantness, change in secondary hyperalgesia) across the whole group, as well as sex disaggregated analyses. For the whole group analysis, STAI-T and BDI were modeled as nuisance covariates (see Results-Questionnaires below). For the disaggregated analysis in males, there were no differences in questionnaire measures, and so no nuisance covariates were modeled. For the sex disaggregated analysis in females, STAI-T was significantly different across intervention arms, and was added as a nuisance covariate. Post-hoc independent t-tests were used to determine differences in outcome measures between stimulation types. Statistical significance was set at p <0.05 adjusted for three outcome measures: p <0.0167.

#### Study 2

Note that as there was only one group for Study 2, we did not perform any statistical comparisons on questionnaires. Descriptive statistics were performed and are presented (see Figure S4). All statistical analyses were performed in SPSS (v.28, IBM, Armonk, NY). There were three key outcome measures in the study: (1) pain intensity, (2) pain unpleasantness and (3) secondary hyperalgesia. To compare these outcome measures between sessions, we performed three paired-sample t-tests (one for each outcome measure). In cases where data were not normally distributed (Shapiro-Wilk’s test), a non-parametric Wilcoxon signed-rank test was used. Statistical significance was set at p <0.05 adjusted for three outcome measures: p <0.0167.

We further performed Bayes factor analysis to determine whether the null hypotheses could be accepted (i.e., no change in pain intensity, pain unpleasantness or secondary hyperalgesia between control and stimulation conditions). Bayes factor analysis allows the assessment of relative evidence in favor of either the null or alternative hypothesis: Bayes Factor (BF)<0.33 is taken as strong evidence of the null hypothesis, and BF>3 is taken as strong evidence in favor of the alternative hypothesis [41]. BFs between 0.33 and 3 are considered to provide no evidence in favor of either hypothesis. The analysis was performed in SPSS v. using the Online Calculator developed by Dienes [12]. To estimate priors, we used the mean difference and the mean standard error from the itbPMS condition from Study 1.

## RESULTS

### Study 1

#### Descriptive Statistics

We compared itbPMS, ctbPMS, TENS, and ctbPMS sham (sham) treatments with four independent groups of participants assigned to one of the different interventions. Figure S1 and Table S1 provide a summary of sample demographic data for Study 1. The itbPMS group consisted of 14 males (mean ± SD age: 25.0 ± 4.10 years) and 12 females (25.1 ± 4.16 years); the ctbPMS group consisted of 14 males (25.7 ± 5.46 years) and 13 females (25.4 ± 5.20 years); the TENS group consisted of 13 males (25.7 ± 5.28 years) and 10 females (26.0 ± 5.01); the sham group consisted of 13 females and 11 males (24.3 ± 5.39 years). Self-reported gender, religious affiliation, and other demographic information can be found summarized in Table S1. Across all arms, the average PHP stimulation temperature used was 45.3 ± 1.5 °C, ranging from 41.0°C - 48.5°C (see Table S2).

#### Questionnaires

We assessed whether there were differences in the self-report mood questionnaires, BDI-II, STAI-S, and STAI-T at baseline. First, there was no significant difference in BDI-II and STAI-S responses between groups (BDI-II: W_3,96_= 2.79, p = 0.050; STAI-S: W_3,96_=2.15, p= 0.105). There was, however, a significant difference between groups in STAI-T baseline characteristics (W_3,96_=3.85, p= 0.015; Table S3).

Sex-disaggregated analysis revealed no significant differences in BDI-II or STAI-S in either males or females. Sex-disaggregated analysis revealed no significant STAI-T differences between males across intervention arms (W_3,48_=2.06, p= 0.130), but did identify a significant difference between females across intervention arms (W_3,44_=3.13, p=0.045). Given that STAI-T was significantly different between intervention arms, it was included as a nuisance covariate all group comparison models. Furthermore, given that BDI was on the cusp of significance (p=0.050), we felt it prudent to include it as a nuisance covariate. Finally, we did not include STAI-S as a nuisance covariate, given it was not significant in any of the models.

#### Group Differences

The pain intensity model (F_3,96_= 1.390, p = 0.251) and pain unpleasantness model (F_3,96_= 0.856, p= 0.467) were not significant, therefore case-wise comparisons were not performed (see Figure 3). However, the secondary hyperalgesia model was significant (F_3,96_= 4.828, p= 0.004; Figure 3). *Post hoc* pairwise comparisons showed significant differences between itbPMS vs TENS (p= 0.00037), ctbPMS vs. TENS (p= 0.011), and TENS vs sham (p= 0.011). In other words, itbPMS, ctbPMS and sham rPMS had significant reductions in the area of secondary hyperalgesia, while TENS had no effect. No other comparisons were significant.

**Figure 3:**
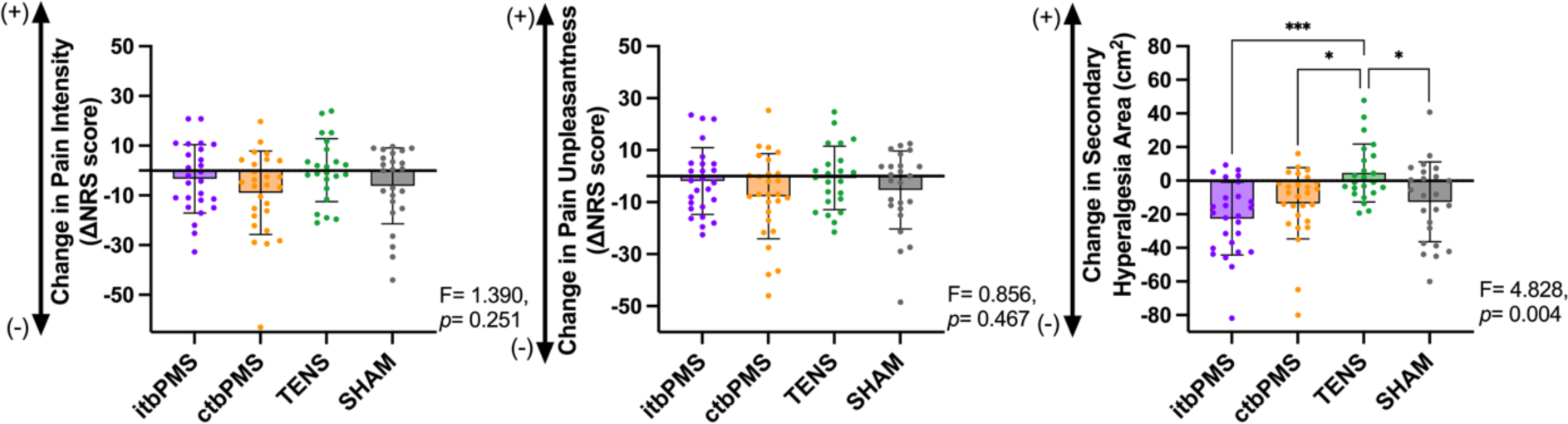
Analysis of intensity, unpleasantness, and secondary hyperalgesia data. The changes in NRS scores were represented as difference in (intervention – control session) for all intervention arms. Significance is shown with stars (*p<0.05, *** p<0.001). No other comparisons were significant.

Additionally, we averaged all subject secondary hyperalgesia maps during the control and treatment sessions to visualize the effects of stimulation for the ITBPMS, ctbPMS, TENS, and sham interventions (see Figure 4).

**Figure 4:**
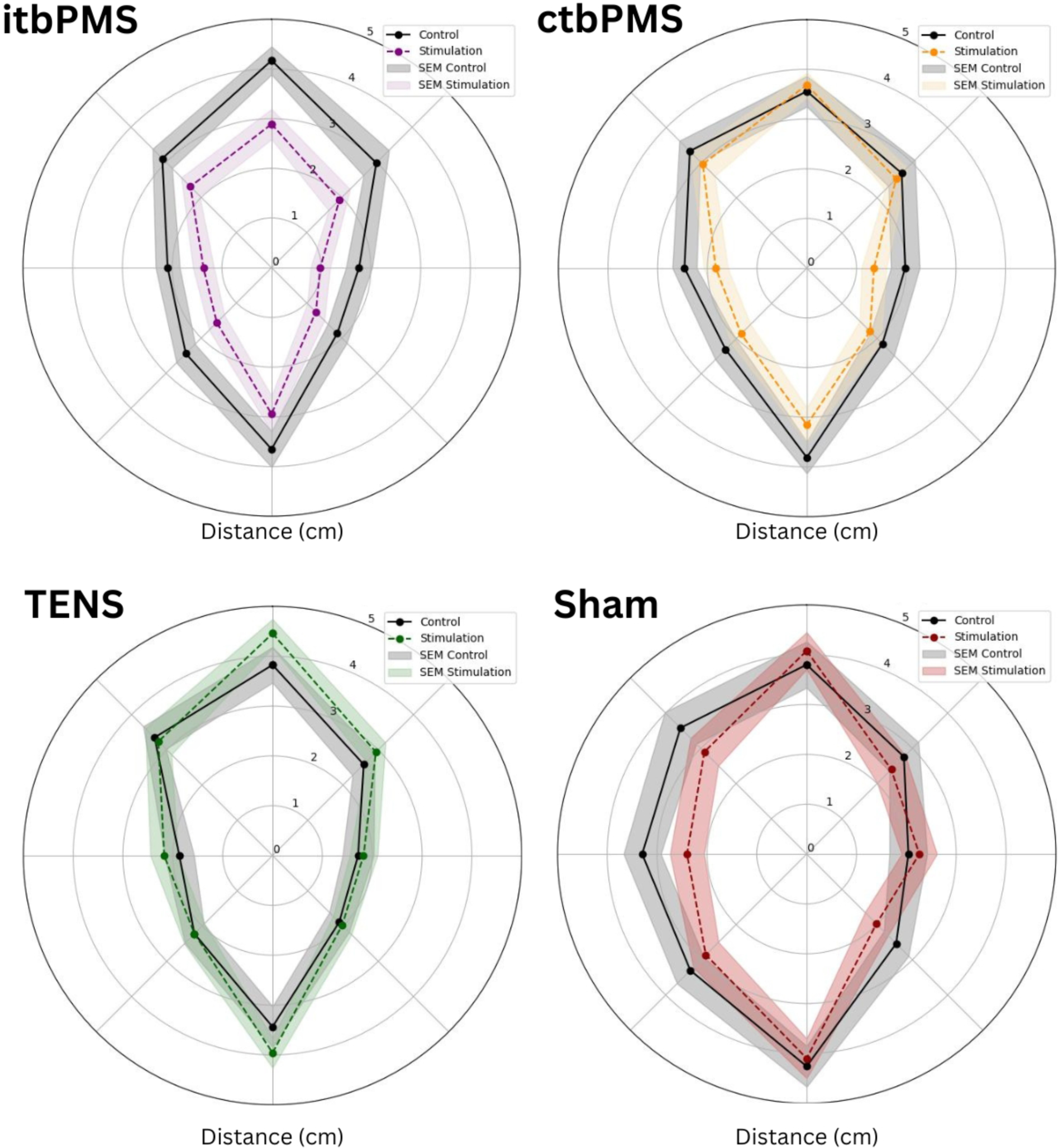
Average secondary hyperalgesia map of all conditions within each intervention arm. The area of secondary hyperalgesia from the intervention session (colored lines) and control session (black) are shown on radial plots in cm. Standard error of the mean is shown in grey for control session, and in color for the stimulation session.

### Study 2

Table S4 summarizes the sample demographics. This study aimed to investigate whether itbPMS masked by pink noise (PN) could significantly reduce a thermally induced secondary hyperalgesia area compared to control condition—i.e., does masking the sound emitted by rPMS abolish the analgesic effect? The study included 24 participants (mean ± SD = 20.7 ± 0.62 years): 12 males (20.7 ± 0.62 years) and 12 females (20.8 ± 0.651 years). Self-reported gender, religious affiliation, and race for each participant can be found summarized in Table S4, and summary scores for mood questionnaires (BDI-II, STAI-S, and STAI-T) are provided in Table S5.

We assessed if there were any differences in reported pain intensity, pain unpleasantness, and secondary hyperalgesia area between control and intervention sessions with paired samples t-tests. We found no significant changes in intensity (t=-0.566, p=0.577) or secondary hyperalgesia (t=-0.498, p=0.310; Figure 5). Additionally, as unpleasantness was non-parametrically distributed, we used the Wilcoxon signed-rank test and found no significant changes (W=187, p=0.304). We further calculated BF for each of the comparisons. We found evidence for the null hypothesis for pain intensity (BF = 0.051, pain unpleasantness (BF=0.043), and secondary hyperalgesia (BF=7.70x10^-8^). In other words, there were no differences in any of the outcome measures between the control and stimulation conditions. Sex-disaggregated analyses also revealed no significant changes in any outcome measures.

**Figure 5:**
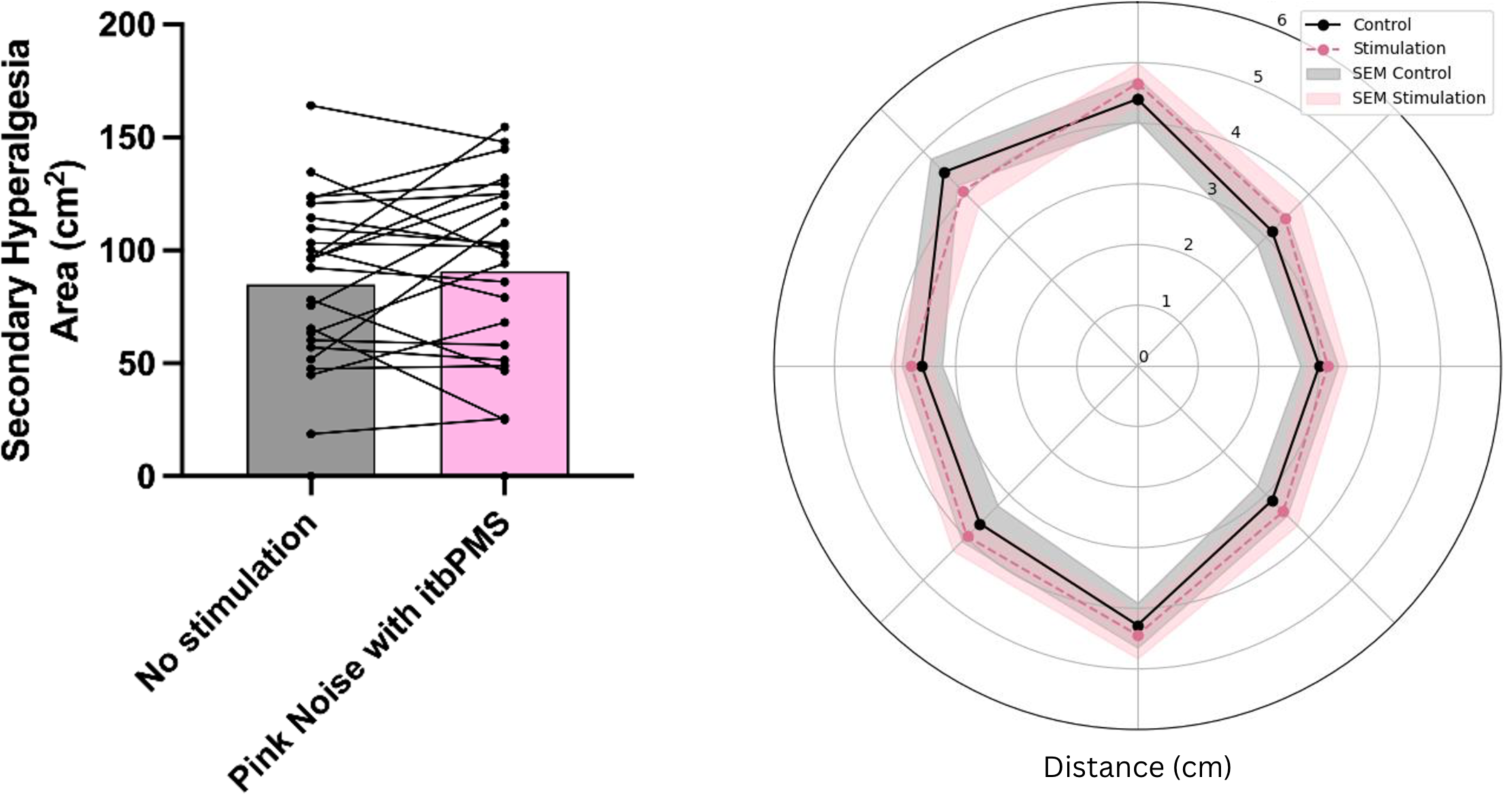
Within-subject comparison of baseline (no stimulation) and itbPMS stimulation with pink noise. No significant changes in secondary hyperalgesia (p=0.501) were reported when participants received no stimulation compared to itbPMS with pink noise. The bar chart (left panel) indicates the mean value of each session, and each line represents a participant. The area of secondary hyperalgesia (right panel) from the intervention session (colored lines) and control session (black) are shown on radial plots in centimeters. Standard error of the mean is shown in grey for control session, and in pink for the itbPMS+pink noise condition.

## DISCUSSION

This study represents the largest controlled investigation of rPMS analgesia for experimental pain to date. We compared the analgesic efficacy of two rPMS stimulation protocols (ctbPMS and itbPMS) against TENS, a peripheral stimulation technique that is commonly used for pain management worldwide. Importantly, we also included a sham rPMS protocol where TMS and PMS naïve participants heard the itbPMS stimulation sound as it was placed over their arm but received no actual magnetic stimulation. We hypothesized that itbPMS and ctbPMS — but not sham — would reduce pain intensity, pain unpleasantness, and secondary hyperalgesia evoked by a phasic heat pain (PHP) paradigm with similar efficacy to TENS. Contrary to our hypothesis, none of the rPMS stimulations or TENS reduced reported pain intensity or unpleasantness. However, ctbPMS and itbPMS significantly reduced the area of secondary hyperalgesia created by the PHP, whereas TENS – which is considered the clinical standard for non-invasive, non-pharmaceutical pain relief — did not. Unexpectedly, sham rPMS, which involved auditory but no peripheral nerve stimulation, also significantly reduced secondary hyperalgesia compared to TENS. This finding suggested that the sham had greater analgesic effectiveness than TENS, a clinically established but inaudible treatment. This led us to design a second study specifically investigating the auditory contributions to rPMS analgesic effects. We discovered that masking the rPMS stimulation sound with pink noise eliminated its modulatory effect on secondary hyperalgesia. Our results suggest that rPMS can reduce experimentally induced secondary hyperalgesia (a marker of central sensitization) and the analgesic properties of rPMS seem to be driven primarily by auditory rather than peripheral nerve stimulation.

Our study included three pain outcome measures: self-reported pain intensity and pain unpleasantness, and the area of secondary hyperalgesia elicited by PHP. In contrast with previous experimental studies of rPMS [29,35,39], we did not see a reduction in pain intensity nor pain unpleasantness following rPMS stimulation. Our key outcome measure was secondary hyperalgesia, a hallmark of central sensitization, where the region around the site of injury becomes more sensitive to noxious stimuli. The area of secondary hyperalgesia is thought to reflect the degree of sensitization of the second order neurons in the spinal cord [21,30,52]. Therefore, we used the size of secondary hyperalgesia area as a measure of how effective each of our five interventions were at preventing or reducing central sensitization, as this could potentially be used as a tool to prevent the transition to persistent and/or chronic pain [32,57]. For example, one study found that the area of secondary hyperalgesia has been associated with the severity of postsurgical pain in individuals undergoing abdominal surgery [14]. They further reported that treatment groups with larger postsurgical areas of secondary hyperalgesia had longer lasting pain up to one year post-surgically [33]. Interestingly, the authors found that pain intensity measurements were not significantly correlated with the reduction in secondary hyperalgesia area, indicating that they are likely capturing different aspects of the pain experience, and have different underlying mechanisms.

Although using measures of secondary hyperalgesia in experimental studies in healthy adults is relatively uncommon, we are not the first to use this outcome to measure therapeutic outcomes. For example, Salomons and colleagues found that a multi-day cognitive behavioral intervention significantly reduced the area of secondary hyperalgesia induced by a series of painful thermal stimuli and was also associated with reduced pain catastrophizing [42]. Similarly, Meeker and colleagues found that anodal motor cortex transcranial direct current stimulation (tDCS) reduced areas of secondary hyperalgesia through modulation of the descending pain modulatory network [37]. Finally, Iannetti and colleague found that gabapentin was associated with reduced activity in the brainstem only when participants experienced experimentally induced central sensitization, measured through the area of secondary hyperalgesia [26]. Together, these studies indicate that secondary hyperalgesia is a modifiable clinically relevant outcome measure, which provides an indirect measure of central sensitization. Such insights can aid in development of targeted therapies aimed at reducing central sensitization and, consequently, chronic pain.

In this study, we used a theta-burst stimulation. Theta oscillations are between 4 and 8Hz, and theta burst stimulation is delivered at 5Hz. Theta rhythms play a key role in memory functions [10,20,23], but their role in pain is less well understood. In chronic pain, theta oscillations have higher power compared to healthy individuals [15,44,56]. Notably, two studies found that patients with intractable neuropathic pain had increased theta power, and after effective treatment with thalamic lesioning, theta oscillations normalized [44,49]. Furthermore, evidence suggests that cortical theta burst stimulation can alleviate diabetic neuropathic pain [50]. However, the location of stimulation (the primary motor cortex or the dorsolateral prefrontal cortex) did not affect outcomes, supporting the concept that the auditory component of the stimulation may be driving the observed effects, in line with our study.

The neural mechanism by which the sound of rPMS reduced central sensitization is unclear. One possibility is that the theta-burst stimulation entrained brain oscillations, i.e., auditory entrainment [31]. Previous studies have shown that entrainment effects can be supramodal [1]—i.e., that entrainment with one modality can affect behaviour associated with another modality. Furthermore, entrainment can affect how sensory inputs are perceived [36]. Most studies investigating brain oscillations and pain have focused on alpha and gamma oscillations as these are often the most prominent oscillation bands found to be associated with pain intensity [9,16-19,22,51,58].

One surprising finding in our study was the ineffectiveness of TENS in reducing pain intensity, pain unpleasantness, or secondary hyperalgesia. Although TENS is often used in clinical pain care, the experimental evidence for its ability to relieve pain varies significantly. Recent meta-analyses [27,54,55] found that TENS reduced pain intensity over placebo TENS, but the results were not consistent across all studies, with significant heterogeneity in the pooled data. The authors concluded that while there is some evidence supporting TENS for acute pain, the quality of evidence is moderate to low due to small sample sizes and methodological limitations.

The relative contributions of peripheral nerve stimulation, distraction by rhythmic auditory stimuli or entrainment, and treatment expectations and placebo to the analgesic effect of rPMS require further investigation. Because rPMS is a non-invasive technique with limited side effects, we believe our sham rPMS condition was effective in simulating the real rPMS condition. Unlike TENS which has direct contact with the skin, the rPMS device in our study does not make physical contact and – depending on the level of stimulation delivered – produced limited perceived sensations of tingling or involuntary flexing of the fingers. These physical indicators of rPMS stimulation were not known to participants as they were naïve to receiving peripheral magnetic stimulation of any kind.

One limitation of the present study is the absence of a measure of participants’ expectations about how the stimulation would impact their pain. Previous studies of experimental pain have shown that an individual’s expectations regarding an intervention can influence their subsequent pain report [5,6,8], and there is robust clinical evidence in the placebo literature that a patient’s expectations about a treatment are correlated with their pain outcomes [11]. However, if these expectations are frequently violated, they have reduced effects [24]. Future studies should include this measure to discern how expectations may modulate analgesic experience. Furthermore, the analgesic efficacy of sham rPMS and the lack of statistically significant analgesia in the pink noise rPMS condition does not preclude some contribution of peripheral nerve stimulation in rPMS analgesia. The stimulation parameters used in our and other rPMS studies are based on standards from cortical stimulation paradigms, and the ideal parameters for peripheral nerve stimulation have yet to be established. Future work should investigate different levels of stimulation with and without the corresponding accompanying auditory stimulation and include electromyography of the targeted peripheral muscles to establish if peripheral stimulation alone is sufficient to induce analgesia and whether the combination of peripheral and auditory stimulation provides more effective relief than either component in isolation.

There are significant therapeutic advantages to auditory theta burst pain management. First, it would be non-invasive, low-cost, accessible and easy to deliver. It would not require expensive magnetic stimulation devices. Second, the selective effects of theta rhythms on reducing or preventing central sensitization, as observed in this study, could be useful as a peri-surgical prophylaxis to prevent post-surgical persistent pain. However, further work is required to establish the reliability and optimization of theta auditory stimulation for acute pain management.

Overall, this study is the first to show that the analgesic properties of rPMS in acute experimental pain may be largely attributed to its auditory component rather than peripheral nerve stimulation. Characterizing the mechanism through which rPMS induces pain relief in other types of pain (clinical, experimental models of neuropathic pain) is an important step before its translation into clinical practice. As rPMS has gained increasing attention as an attractive alternative to current pharmacological treatments because its non-invasive and has few side effects, more high-quality controlled studies are needed to disentangle the mechanisms underlying rPMS-induced pain relief. Indeed, if auditory theta-range stimuli are effective at reducing central sensitization, then the study of underlying mechanisms warrants further investigation, as it presents a low-cost, easy pain management solution.

## Data availability statement

Data are included in the supplement of the manuscript.

## Statement of Contributions

EEO, NO, SSA: Study design, data collection, analysis, writing

RT, NWDV: data collection, editing

JSK, LH, AF, AM, DAS: Study design, review, editing

MM: Conception of study, funding, analysis, editing, approval of final manuscript

## ACKNOWLEDGMENTS

EE Osokin was funded by a Medical Institute of Berezina Sergey Scholarship. M Moayedi is supported by a University of Toronto Centre for the Study of Pain — Pain Scientist Award, a Canada Research Chair (Tier 2) in Pain Neuroimaging, and the Bertha Rosenstadt Endowment Fund at the Faculty of Dentistry, University of Toronto. The authors have no conflicts to report. This study was funded through Moayedi’s discretionary funds. SS Abssy is supported by University of Toronto Excellence Award.

Dr. Liat Honigman, who is a co-author on this paper, was a beloved lab member and research associate in the Centre for Multimodal Sensorimotor and Pain Research (CMSPR) at the University of Toronto. She died on March 28, 2024. This, and many other projects in the lab would not have been possible without her expertise, guidance, and support. Liat was kind, smart, and a mentor to many trainees in the CMSPR. She held the highest standards for science, but believed in a compassionate approach to research. She is truly missed.

