## Supplementary figures and images for "It’s the Sound, not the Pulse: Peripheral Magnetic Stimulation Reduces Central Sensitization through Auditory Modulatory Effects"

### Figure S1

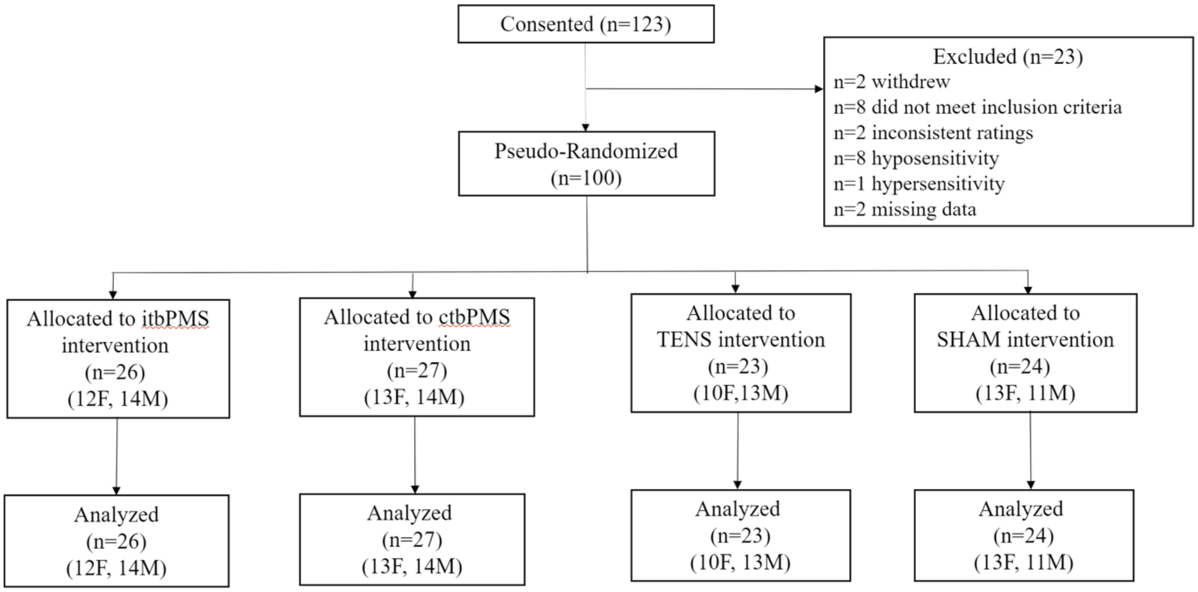


**Figure S1. Consolidated Standards of Reporting Trials (CONSORT) flow diagram for Study 1**.

### Figure S2

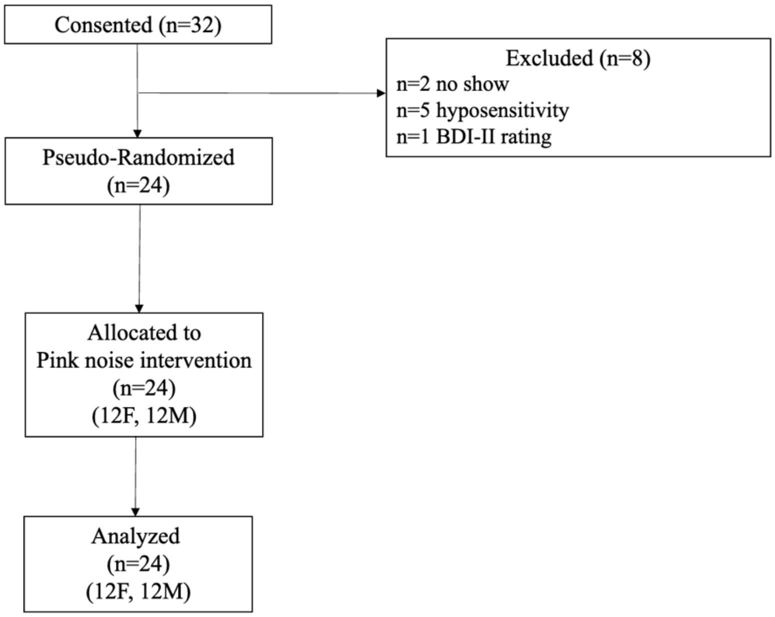


**Figure S2. Consolidated Standards of Reporting Trials flow diagram for Study 2.**
