## Supplementary material for "It’s the Sound, not the Pulse: Peripheral Magnetic Stimulation Reduces Central Sensitization through Auditory Modulatory Effects": Table S1

**Table S1: Demographic information and of participants in Study 1.**

| | | Intervention Arm | | |
|---|---|---|---|---|
| | itbPMS | ctbPMS | TENS | Sham |
| Age *(mean ± SD years)* | 25.0 ± 4.1 | 25.4 ± 5.2 | 26 ± 5.01 | 24.3 ± 5.39 |
| Race (# individuals) | | | | |
| East Asian | 8 | 8 | 6 | 4 |
| South Asian | 5 | 3 | 2 | 4 |
| South East Asian | 1 | 0 | 1 | 2 |
| Latin American | 4 | 2 | 2 | 1 |
| Middle Eastern | 2 | 1 | 2 | 0 |
| Mixed Heritage | 1 | 1 | 1 | 1 |
| White/European | 4 | 10 | 8 | 8 |
| Black Caribbean | 0 | 1 | 0 | 1 |
| Black African | 0 | 0 | 0 | 3 |
| Do Not Know/Prefer not to answer | 0 | 1 | 1 | 0 |
| Religious/Spiritual Affiliation | | | | |
| Atheism | 6 | 2 | 2 | 0 |
| Buddhism | 0 | 1 | 1 | 0 |
| Christianity | 0 | 6 | 3 | 4 |
| Hinduism | 3 | 0 | 0 | 1 |
| Islam | 1 | 4 | 2 | 1 |
| Jainism | 0 | 0 | 1 | 0 |
| Judaism | 2 | 4 | 0 | 0 |
| Spiritual | 1 | 2 | 0 | 2 |
| Not Religious/Spiritual | 12 | 8 | 11 | 15 |
| Other | 0 | 0 | 1 | 0 |
| Prefer not to answer | 1 | 0 | 2 | 1 |
| Sex assigned at birth | | | | |
| Female | 12 | 13 | 10 | 13 |
| Male | 14 | 14 | 13 | 11 |
| Gender | | | | |
| Man | 14 | 13 | 13 | 11 |
| Woman | 12 | 13 | 10 | 13 |
| Non-Binary | 0 | 0 | 0 | 0 |
| Trans | 0 | 1 | 0 | 0 |
