## Supplementary material for "It’s the Sound, not the Pulse: Peripheral Magnetic Stimulation Reduces Central Sensitization through Auditory Modulatory Effects": Table S2

**Table S2. Stimulus temperatures for Study 1.**

| | | Intervention Arm | | | | |
|---|---|---|---|---|---|---|
| | | itbPMS | ctbPMS | TENS | Sham | All Arms |
| PHP Temperature | | | | | | |
| | Males | 45.6 ± 1.6 | 45.4 ± 1.1 | 45.5 ± 0.9 | 45.4 ± 2.1 | 45.5 ± 1.4 |
| | Females | 44.8 ± 1.5 | 45.2 ± 1.4 | 44.7 ± 1.7 | 45.5 ± 1.9 | 45.1 ± 1.6 |
| | All subjects | 45.2 ± 1.6 | 45.3 ± 1.2 | 45.1 ± 1.3 | 45.4 ± 2.0 | 45.3 ± 1.5 |
| PHP Temperature Range | | | | | | |
| | Males | 42.0 - 47.0 | 43.0 - 47.0 | 44.0 - 47.0 | 42.0 - 48.5 | 42.0 - 48.5 |
| | Females | 41.0 - 47.0 | 42.0 - 47.0 | 42.0 - 46.0 | 41.0 - 47.5 | 41.0 - 47.5 |
| | All subjects | 41.0 - 47.0 | 42.0 - 47.0 | 42.0 - 47.0 | 41.0 - 48.5 | 41.0 - 48.5 |
