## Supplementary material for "It’s the Sound, not the Pulse: Peripheral Magnetic Stimulation Reduces Central Sensitization through Auditory Modulatory Effects": Table S3

**Table S3: Group differences in mood questionnaires for Study 1.**

| | | itbPMS | ctbPMS | TENS | Sham | Anova F (p) |
|---|---|---|---|---|---|---|
| State-Trait Anxiety Inventory | | | | | | |
| State | Males | 29.3 ± 5.67 | 34.0 ± 9.20 | 30.1 ± 8.72 | 35.1 ± 8.28 | 1.68 (0.196) |
| | Females | 24.9 ± 3.37 | 26.6 ± 4.43 | 27.9 ± 4.13 | 29.2 ± 7.08 | 1.69 (0.196) |
| | All Subjects | 27.3 ± 5.16 | 30.4 ± 8.09 | 29.1 ± 7.11 | 31.8 ± 8.38 | 2.15 (0.105) |
| Trait | Males | 33.1 ± 6.75 | 36.1 ± 8.55 | 39.2 ± 9.48 | 39.7 ± 7.56 | 2.06 (0.130) |
| | Females | 27.8 ± 5.89 | 35.4 ± 7.34 | 33.6 ± 6.8 | 34.4 ± 10.3 | 3.13 (0.045) |
| | All Subjects | 30.7 ± 6.95 | 35.8 ± 7.85 | 36.8± 8.84 | 36.8 ± 9.73 | 3.85 (0.015) |
| Beck's Depression Index II | | | | | | |
| | Males | 2.6 ± 2.45 | 5.6 ± 5.19 | 5.3 ± 6.48 | 4.2 ± 3.79 | 1.62 (0.209) |
| | Females | 2.7 ± 2.43 | 5.3 ± 4.91 | 2.5 ± 2.97 | 3.7 ± 3.95 | 1.55 (0.229) |
| | All Subjects | 2.6 ± 2.45 | 4.4 ± 4.99 | 4.5 ± 5.39 | 4.2 ± 3.96 | 2.79 (0.050) |

The State-Trait Anxiety Inventory (STAI) and Beck's Depression Inventory II (BDI-II) were collected before the experimental portion of the study. The mean scores ± SD were calculated for each intervention arm and within each sex. Results of a one-way ANOVA are provided ($\alpha$=0.05).
