## Supplementary material for "It’s the Sound, not the Pulse: Peripheral Magnetic Stimulation Reduces Central Sensitization through Auditory Modulatory Effects": Table S4

**Table S4. Demographic information and descriptive statistics of participants in Study 2**

| | | |
|---|---|---|
| Age *(mean ± SD years)* | | 20.7± 0.62 |
| Race (# individuals) | | |
| | East Asian | 7 |
| | South Asian | 3 |
| | South East Asian | 2 |
| | Latin American | 1 |
| | Middle Eastern | 0 |
| | Mixed Heritage | 0 |
| | White/European | 8 |
| | Black Caribbean | 0 |
| | Black African | 2 |
| | Do Not Know/Prefer not to answer | 1 |
| Religious/Spiritual Affiliation | | |
| | Atheism | 1 |
| | Buddhism | 0 |
| | Christianity | 7 |
| | Hinduism | 1 |
| | Islam | 3 |
| | Jainism | 0 |
| | Judaism | 0 |
| | Spiritual | 1 |
| | Not Religious/Spiritual | 11 |
| | Other | 0 |
| | Prefer not to answer | 0 |
| Sex assigned at birth | | |
| | Female | 12 |
| | Male | 12 |
| Gender | | |
| | Man | 12 |
| | Woman | 12 |
| | Non-Binary | 0 |
| | Trans | 0 |
