## Supplementary material for "It’s the Sound, not the Pulse: Peripheral Magnetic Stimulation Reduces Central Sensitization through Auditory Modulatory Effects": Table S5

**Table S5: Study 2 Participant Questionnaire Data.**

| | Males | Females | All Subjects |
|---|---|---|---|
| State-Trait Anxiety Inventory | | | |
| State | 31.8 ± 11.3 | 35.4± 11.1 | 33.6 ± 11.1 |
| Trait | 41.3 ± 10.0 | 39.8 ± 9.84 | 40.6 ± 9.76 |
| Beck's Depression Index II | | | |
| | 9.41 ± 7.91 | 7.08 ± 4.93 | 8.25 ± 6.56 |

The State-Trait Anxiety Inventory (STAI) and Beck's Depression Inventory II (BDI-II) were collected before the experimental portion of the study. The mean scores ± SD were calculated.
